# Identification of Platform-Independent Diagnostic Biomarker Panel for Hepatocellular Carcinoma using Large-scale Transcriptomics Data

**DOI:** 10.1101/758250

**Authors:** Harpreet Kaur, Anjali Dhall, Rajesh Kumar, Gajendra P. S. Raghava

## Abstract

The high mortality rate of hepatocellular carcinoma (HCC) is primarily due to its late diagnosis. In the past numerous attempts have been made to design genetic biomarkers for the identification of HCC; unfortunately, most of the studies are based on a small dataset obtained from a specific platform or lack of their reasonable validation performance on the external datasets. In order to identify a universal expression-based diagnostic biomarker panel for HCC that can be applicable across multiple platforms; we have employed large scale transcriptomic profiling datasets containing a total of 2,306 HCC and 1,655 non-tumorous tissue samples. These samples were obtained from 29 studies generated by mainly four types of profiling techniques include Affymetrix, Illumina, Agilent and High-throughput-seq, which implemented a wide range of platforms. Firstly, we scrutinized 26 genes that are differentially expressed or regulated in uniform pattern among numerous datasets. Subsequently, we identified three genes (*FCN3, CLEC1B*, & *PRC1)* panel-based HCC biomarker using different machine learning techniques include Simple-threshold based approach, Extra Trees, Support Vector Machine, Random Forest, K Neighbors Classifier, Logistic Regression *etc*. Three-genes panel-based HCC biomarker classified HCC samples and non-tumorous samples of training and three external validation datasets with an accuracy between 93 to 98% and AUROC (Area Under Receiver Operating Characteristic curve) in a range of 0.97 to 1.0. Furthermore, the prognostic potential of these genes was evaluated on TCGA cohort and GSE14520 cohort using univariate survival analysis revealed that these genes are independent prognostic indicators for various types of the survivals, *i.e.* OS (Overall Survival), PFS (Progression-Free Survival), DFS/RFS (Disease-Free Survival/Recurrence-Free Survival) and DSS (Disease-Specific Survival) of HCC patients and significantly stratify high-risk and low-risk HCC patients (p-value <0.05). In conclusion, we identified a universal platform-independent three genes-based biomarker that can predict HCC patients with high precision; also possess significant prognostic potential. Eventually, to provide service to the scientific community, we developed a webserver HCCPred based on the above study (http://webs.iiitd.edu.in/raghava/hccpred/).

## 1 Introduction

Cancer is a heterogeneous disease driven by genomic and epigenomic changes within the cell (Chatterjee et al., 2018; Dawson and Kouzarides, 2012; Flavahan et al., 2017; Kagohara et al., 2018; Kamel and Al-Amodi, 2017; Kumar et al., 2019; Nagpal et al., 2015; Narrandes and Xu, 2018; Nebbioso et al., 2018; Sharma et al., 2010). Gene dysregulation is considered a hallmark of cancer. Among the 22 common cancer type, Hepatocellular carcinoma (HCC) ranks at sixth in terms of frequency of occurrence and fourth at cancer-related mortality (Siegel et al., 2019). The aetiology of HCC can be induced by multiple factors, especially virus infection, alcoholic cirrhosis and consumption of aflatoxin-contaminated foods (Ho et al., 2016). Although various traditional and locoregional treatment strategies such as Hepatic resection (RES), Percutaneous ethanol injection (PEI), radiofrequency ablation (RFA), microwave ablation (MWA) and trans-arterial chemotherapy infusion (TACI) have improved the survival rate but patient with HCC still have a late diagnosis and poor prognosis (Tian et al., 2018).

In the past, several studies focus on the identification of biomarkers by comparing the global gene expression changes between cancer tissue and non-tumorous tissues (Cai et al., 2017, 2019; Emma et al., 2016; Gao et al., 2015; Jia et al., 2007; Jiao et al., 2019; Kang et al., 2015; Komatsu et al., 2016; Li et al., 2017, 2018; Liao et al., 2018; Liu et al., 2015; Marshall et al., 2013; Meng et al., 2018; Shirota et al., 2001; Wang et al., 2018; Xia et al., 2019; Xu et al., 2018; Zhang et al., 2017, 2019; Zheng et al., 2018). Such analyses yield hundreds or thousands of gene signature that are differentially expressed in cancer tissue compared to normal tissue. Thus, making it difficult to identify a universal subset of genes that play a crucial role in neoplastic transformation and progression (Rhodes et al., 2004). The lack of concordance of signature genes among different studies and extensive molecular variation between the patient’s samples restrains the establishment of the robust biomarkers, promising targets and their experimental validation in clinical trials (Vasudevan et al., 2018). The transcriptome signatures have yet to be translated into a clinically useful biomarker, that may be due to a lack of their satisfactory validation performance on independent patient’s cohort.

In this regarding, treatment of HCC remains unsatisfying as only diagnostic and prognostic biomarkers alpha-fetoprotein (AFP) has been established so far. Several other biomarkers AFP-L3, osteopontin, glypican-3 are currently being under investigation for the early diagnosis of HCC patients (Ocker, 2018). Advancement in the genomics has created rich public repositories of microarray and high throughput datasets from numerous studies such as TCGA(The Cancer Genome Atlas Program - National Cancer Institute), GDC(GDC) and GEO(Home - GEO - NCBI), provide the opportunity to study the various aspects of cancer. Thus, novel methods exploring computational approach by merging multiple datasets from different platforms could provide a new way to establish a robust and universal biomarker for disease diagnosis and prognosis with increased precision and reproducibility. Recently, multiple datasets integration approach implemented for identification of robust biomarker panel for pancreatic ductal adenocarcinoma (PDAC) (Bhasin et al., 2016; Klett et al., 2018). Although, various studies employed large-scale data or meta-analysis approaches to identify protein and miRNA expression-based biomarker for HCC diagnosis (Chen et al., 2018b; Ding et al., 2017; Ji et al., 2016, 2018). But, best of our knowledge, RNA-expression data is not explored in this regard for identification of the robust biomarker for HCC diagnosis and prognosis.

To overcome the limitations of existing methods, we made a systematic attempt to identify genetic biomarkers for HCC diagnosis that apply to a wide range of platforms and profiling techniques. One of the objectives of this study is to identify robust gene expression signature for discrimination of HCC samples by the integration of multiple transcriptomic datasets from various platforms. Here, we have collected and analysed a total of 3,961 samples from published datasets, out of which 2,306 and 1,655 are of HCC and normal or non-tumorous tissue samples, respectively. From this, we identified 26 genes, which are commonly differentially expressed in uniform patterns among most of the datasets, which provides a universally activated transcriptional signature of HCC cancer type. Further, we have established a robust “Three-genes based HCC biomarker” implementing different machine learning techniques to distinguish HCC and non-tumorous samples with high precision. Additionally, the survival analysis of HCC patient’s cohorts using these genes revealed their significant prognostic potential in the stratification of high-risk and low-risk patient’s groups. To best of our knowledge, this is the first study in the regards of HCC cancer type, which utilizes such a large-scale transcriptomic dataset from different platforms, for the identification of universal platform-independent diagnostic biomarkers implementing machine learning approaches.

## 2. Materials and Methods

### 2.1 Dataset Collection

#### 2.1.1 Collection of gene expression datasets of HCC

In order to analyse the large-scale gene expression profiles in Hepatocellular carcinoma (HCC), supplementary data for twenty-nine transcriptome studies (GSE102079 (Chiyonobu et al., 2018), GSE22405, GSE98383 (Diaz et al., 2018), GSE84402 (Wang et al., 2017), GSE64041 (Makowska et al., 2016), GSE69715 (Sekhar et al., 2018), GSE51401, GSE62232 (Schulze et al., 2015), GSE45267 (Chen et al., 2018a), GSE32879 (Oishi et al., 2012), GSE19665 (Deng et al., 2010), GSE107170 (Diaz et al., 2018), GSE76427 (Grinchuk et al., 2018), GSE39791 (Kim et al., 2014), GSE57957 (Mah et al., 2014), GSE87630 (Woo et al., 2017), GSE46408, GSE57555 (Murakami et al., 2015), GSE54236 (Villa et al., 2016; Zubiete-Franco et al., 2019), GSE65484 (Dong et al., 2015), GSE31370 (Seok et al., 2012), GSE84598, GSE89377, GSE29721 (Stefanska et al., 2011), GSE14323 (Mas et al., 2009), GSE25097 (Lamb et al., 2011; Tung et al., 2011; Wong et al., 2016), GSE14520 (Roessler et al., 2010; Zhao et al., 2015), GSE36376 (Lim et al., 2013) and TCGA-LIHC), each contains minimum 10 samples. All these datasets were initially obtained using GEOquery package (Bioconductor - GEOquery) from GEO and TCGA-LIHC RNA-seq data with metadata and clinical files was downloaded using gdc-client from GDC data portal, respectively. Then, we manually curated these datasets to ensure that the data contains only expression data from human HCC tissues. For each of the 29 datasets included in the current study, we reviewed the samples profiled. Twenty-nine studies had at least ten samples corresponding to both classes, *i.e.* HCC and normal (or adjacent non-tumor) were considered for the analysis of interest. Further, each dataset carefully investigated to check proper sample ID, type of sample and to remove any irrelevant error in dataset file. Besides, Gene symbol mapped to probe IDs were extracted from respective Platform file (wherever available) and incorporated in the dataset matrix for each dataset.

#### 2.1.2 Pre-processing of datasets

Each retrieved raw dataset (supplementary data) was subjected to a detailed curation process. We have pre-processed each dataset matrix individually from each profiling technique for each platform in a standardized manner. For Affymetrix datasets, raw data files were pre-processed with background correction and eventually RMA values calculated using the Oligo package (Carvalho and Irizarry, 2010). For Illumina datasets, raw files were processed using Limma and Lumi packages (Du et al., 2008; Ritchie et al., 2015) and finally log2 values calculated using in-house R scripts. Similarly, raw Agilent-1-color and Agilent-2-color files were pre-processed using Limma package individually, then A-values were generated, which were further transformed to log2 values. Eventually, the average of multiple probes computed that correspond to a single gene for each dataset individually employing in-house R scripts. TCGA-LIHC dataset contains FPKM values, which were further converted to log2 values. Entrez transcript IDs are mapped to the Gene symbols using Gencode v22.

#### 2.1.3 Datasets used for the Identification of Differentially Expressed Genes (DEG)

Twenty-five out of twenty-nine datasets selected, each containing samples more than ten for initial analysis to identify differentially expressed genes between HCC and normal as given in Figure 1A. These datasets were further analyzed to assign the samples to three different classes, *i.e.*, HCC, adjacent non-tumor and healthy normal samples. Eventually, we generated a total of 27 datasets; twenty datasets contain HCC v/s adjacent non-tumor samples and seven datasets contain HCC v/s healthy normal samples. These datasets encompass 1199 HCC samples and 949 normal or adjacent non-tumor samples.

**Figure1:**
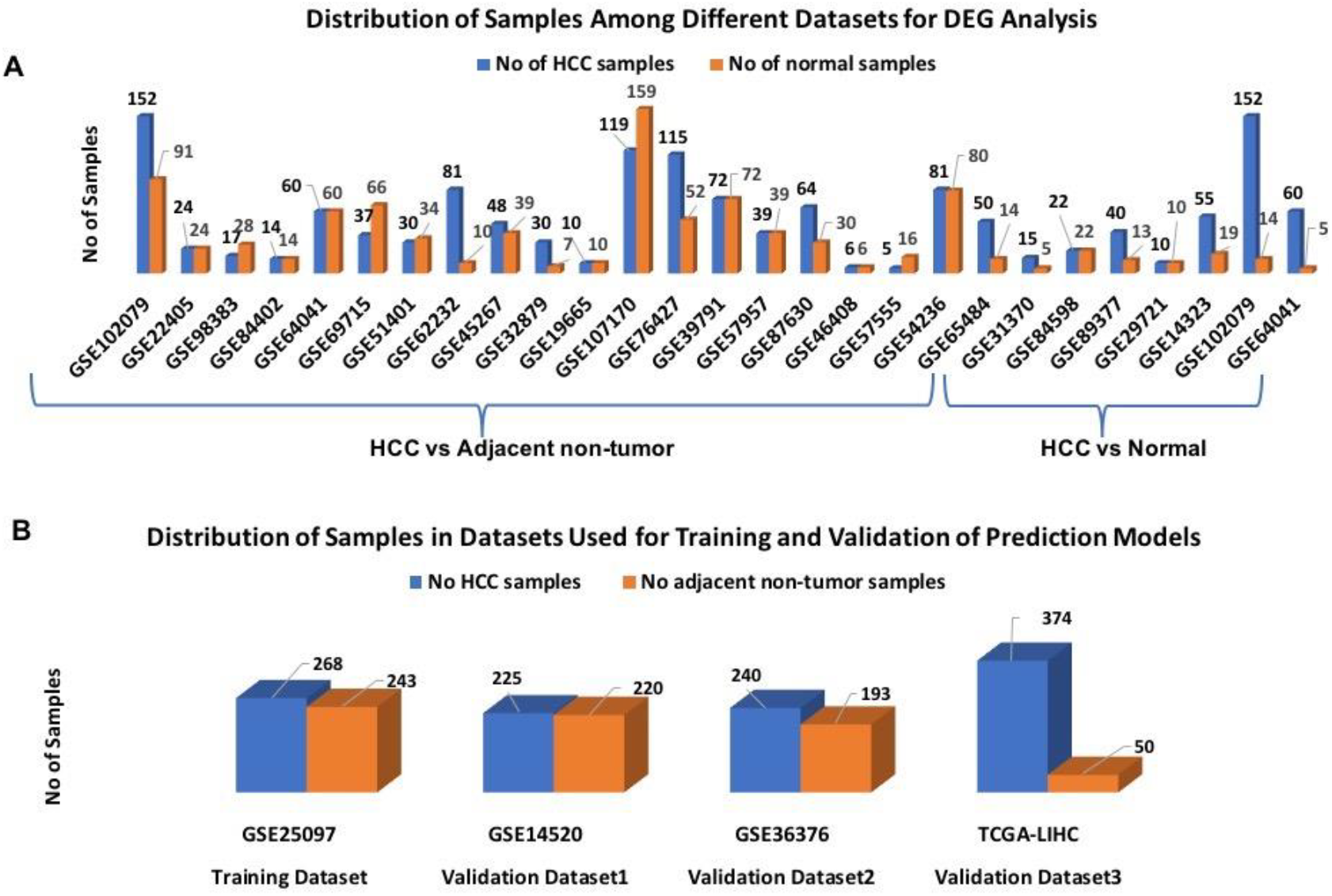
Distribution of Samples among Datasets used in the study

#### 2.1.4 Training and Independent Validation Datasets

Four out of twenty-nine datasets (GSE25097, GSE14520, GSE36376 and TCGA-LIHC datasets) selected each having more than 400 samples, for training the model and validation as given in Figure 1B. To reduce the cross-platform artifacts, quantile normalization is performed using the PreprocessCore package of Bioconductor (GitHub - bmbolstad/preprocessCore) for each dataset and for each profiling technique by keeping training dataset as model (target) matrix. These datasets contain a total of 1107 HCC and 706 adjacent non-tumor samples.

### 2.2 Analysis for the Identification of Differentially Expressed Genes (DEGs)

For each of the 32 microarray datasets, each gene was analyzed for differential expression with Student’s *t-*test (Welch *t*-test and Wilcoxon *t*-test) using in-house Rscript after the assignment of samples to the respective class, *i.e.* cancer or normal. *T*-tests were conducted as two-sided for differential expression analysis. Only those set of genes chosen to define differential expressed genes from each of datasets that are statistically differentially expressed between two classes with Bonferroni adjusted p-value less than 0.01. With aim to identify a selected set of differential expression signatures or “core genes of Hepatocellular carcinoma”, DEGs were compared among all 27 datasets.

### 2.3 Identification of Robust Biomarkers for HCC Diagnosis

#### 2.3.1 Reduction of Features

With intent to reduce the number of genes from the selected the set of signature, *i.e.* “the core genes of Hepatocellular carcinoma”, we used two machine learning feature selection techniques include simple threshold-based approach (Bhalla et al., 2017) and ten-fold cross-validation technique using training cohort (GSE25097). Firstly, we employed a simple threshold-based technique in which we developed single feature-based models to distinguish cancer and normal samples. Single feature-based models are also called threshold-based models wherein genes with a score above the threshold are assigned to cancer class if it is found to be upregulated in cancer and otherwise normal; whereas sample is assigned to normal class if the gene is downregulated in cancerous condition as reported in our previous studies(Bhalla et al., 2017; Kaur et al., 2018). We compute the performance of each model based on a given feature and identify the top 10 features having the highest performance in terms of different performance measures like accuracy, MCC and AUROC, etc.

Further, ten-fold cross-validation technique is implemented to select only most important features from above top 10 selected features (genes) based on their individual performance in terms of all the performance measures (sensitivity, specificity, accuracy, MCC and AUROC) to discriminate cancer and normal samples.

#### 2.3.2 Development of Prediction models implementation Machine Learning Techniques

Here, we have developed the prediction models to distinguish HCC and normal samples based on selected genomic features using various classifiers implementing scikit-learn package. These classifiers include ExtraTrees, Naive Bayes, KNN, Random forest, Logistic Regression (LR), and SVC-RBF (radial basis function) kernel were implemented employing scikit-learn package (Scikit-learn: Machine Learning in Python). The optimization of the parameters for the various classifiers was done by using a grid search with AUROC curve as scoring performance measure for selecting the best parameter.

### 2.4 Performance Evaluation of Models

In the current study, both internal and external validation techniques were employed to evaluate the performance of models. First, the training dataset is used to develop prediction models and ten-fold cross-validation is used for performing internal validation. It is important to evaluate the realistic performance of the model on external validation dataset, which should not be used for training and testing during model development. Therefore, we also implemented external validation. Towards this, we evaluated the performance of our model on three independent gene expression cohorts include GSE14520, GSE36376, and TCGA-LIHC obtained from GEO and The Cancer Genome Atlas (TCGA), each containing more than 400 samples (Supplementary Information File 2, Figure S1C), were not used for training. In order to measure the performance of models, we used standard parameters. Both threshold-dependent and threshold-independent parameters were employed to measure the performance. In the case of threshold-dependent parameters, we measure sensitivity, specificity, accuracy and Matthew’s correlation coefficient (MCC) using the following equations.

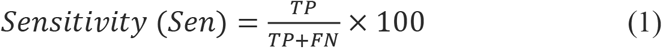

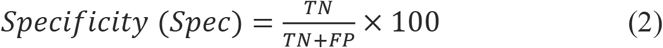

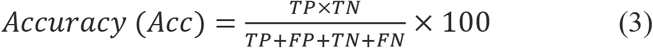

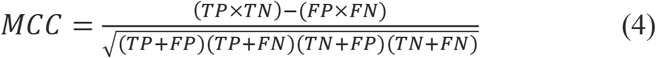

Where, FP, FN, TP and TN are false positive, false negative true positive and true negative predictions, respectively.

While, for threshold-independent measures, we used a standard parameter Area under the Receiver Operating Characteristic curve (AUROC). The AUROC curve is generated by plotting sensitivity or true positive rate against the false positive rate (1-specificity) at various thresholds. Finally, the area under the curve calculated to compute a single parameter called AUROC.

### 2.5 Prognostic Potential of Identified HCC Diagnostic Biomarkers

The prognostic potential of the ***“Three-genes HCC Biomarker”*** was analyzed using gene expression data of TCGA-LIHC and GSE14520 cohorts. The TCGA dataset and GSE14520 dataset contain of 374 and 219 tumor samples with their clinical information extracted from GEO, TCGA and literature (Liu et al., 2018; Roessler et al., 2010), respectively. The clinical characteristics of patients are given in Table S11 (Supplementary Information File 1). The survival univariate analyses and risk assessments were performed by Survival package in R (Therneau). The survival signatures of biomarkers were evaluated by Kaplan-Meier plots, and a log-rank p-value < 0.05 was considered the cut-off to describe the statistical significance in all survival analyses. Here, we analyzed four types of survivals, *i.e.* OS (Overall Survival), DSS (Disease-Specific Survival), DFS (Disease-Free Survival) and PFS (Progression-Free Survival) for TCGA cohort, and two types of survivals, *i.e.* OS and RFS (Recurrence-Free Survival) [also called as Disease-Free Survival (DFS)] for GSE14520 cohort.

### 2.6 Functional Annotation of Signature Genomic Markers

In order to discern the biological relevance of the signature genes, enrichment analysis is performed using Enrichr (Kuleshov et al., 2016). Enrichr executes Fisher exact test to identify enrichment score. It provides Z-score and adjusted p-value, which is derived by applying correction on a Fisher Exact test.

## 3 Results

### Overview

The pipeline of our analysis is illustrated in Figure 2. The details of each step are described below.

**Figure2:**
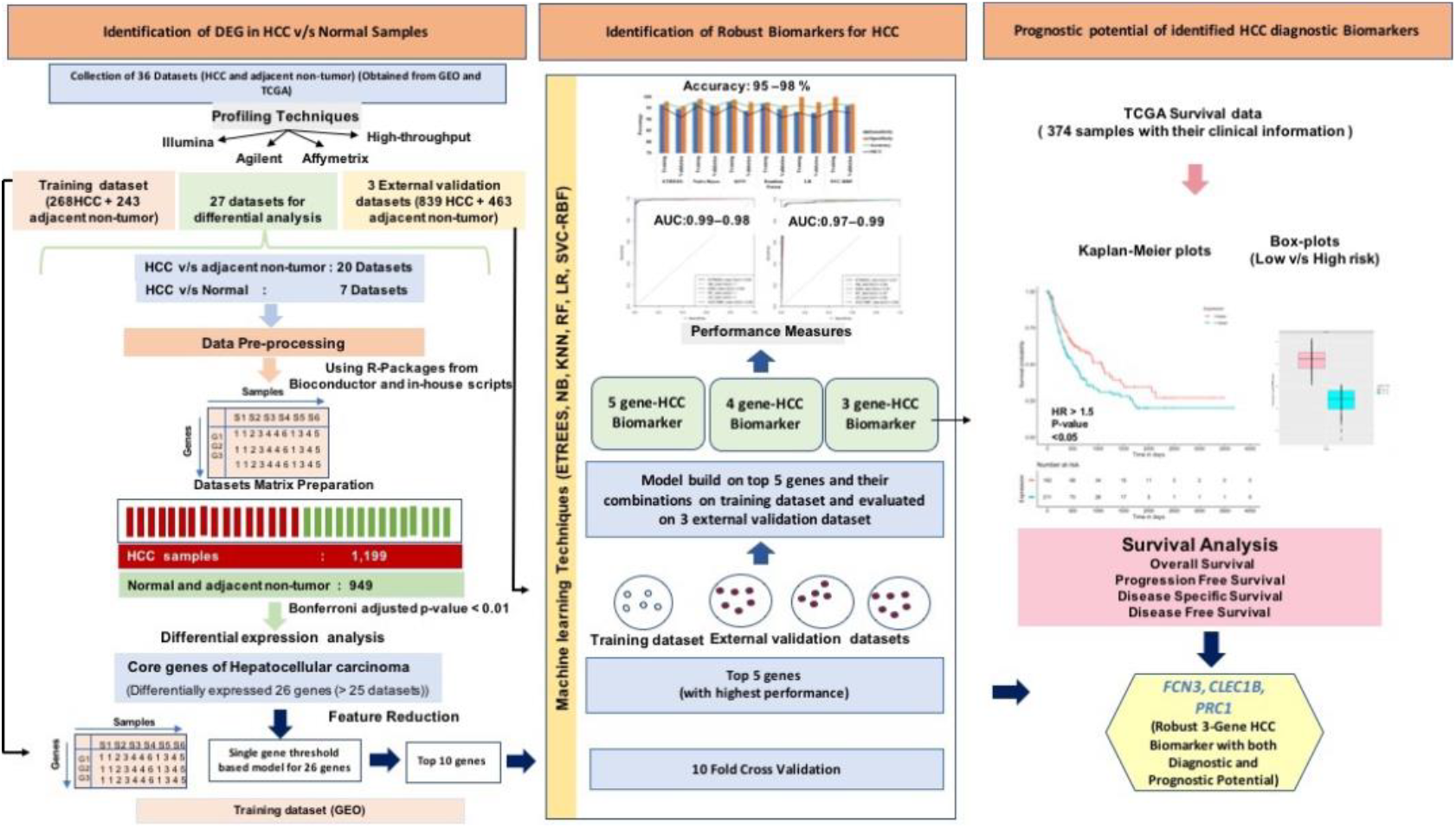
Overview for the analysis implemented in the study

#### 3.1 The Transcriptomic Cores for Hepatocellular Carcinoma

The individual statistical differential expression analyses of 27 gene expression datasets resulted in the identification of hundreds of upregulated and downregulated Differentially Expressed Genes (DEGs) (Supplementary Figure 1). The 9,954 genes are present among each of 27 datasets (Supplementary Information File 1, Table S1). Further, the comparative analysis among all 27 datasets scrutinized 26 genes that are found to statistically differentially expressed in at least more than 80% of the datasets, i.e. 22 datasets. We called these genes as “Core genes for Hepatocellular carcinoma”. Among these 26 genes, 12 are downregulated and 14 are upregulated in HCC in comparison to normal samples. The regulatory patterns of the core genes were consistent among most of the datasets (Table 1). Additionally, the expression pattern of these genes in training and three external validation datasets is shown in Figure S2 (Supplementary Information File 2).

**Table1:** List of genes that are differentially expressed between HCC and adjacent normal or adjacent non-tumor samples with Bonferroni p-values < 0.01.

| Gene | #Up | #Down | #Sig | #Non-sig | Up (%) | Down (%) | Sig (%) | Regulation |
| --- | --- | --- | --- | --- | --- | --- | --- | --- |
| *FCN3* | 1 | 26 | 24 | 3 | 3.70 | 96.30 | 88.89 | Down |
| *CLEC4M* | 2 | 25 | 24 | 3 | 7.41 | 92.59 | 88.89 | Down |
| *FCN2* | 2 | 25 | 24 | 3 | 7.41 | 92.59 | 88.89 | Down |
| *MARCO* | 3 | 24 | 22 | 5 | 11.11 | 88.89 | 81.48 | Down |
| *CRHBP* | 2 | 25 | 22 | 5 | 7.41 | 92.59 | 81.48 | Down |
| *CFP* | 2 | 25 | 22 | 5 | 7.41 | 92.59 | 81.48 | Down |
| **STEAP3** | 2 | 25 | 25 | 2 | 7.41 | 92.59 | 92.59 | Down |
| **HGFAC** | 4 | 23 | 22 | 5 | 14.81 | 85.19 | 81.48 | Down |
| **CLEC1B** | 2 | 25 | 23 | 4 | 7.41 | 92.59 | 85.19 | Down |
| **CXCL12** | 3 | 24 | 24 | 3 | 11.11 | 88.89 | 88.89 | Down |
| **MT1E** | 3 | 24 | 24 | 3 | 11.11 | 88.89 | 88.89 | Down |
| **NSUN5** | 25 | 2 | 24 | 3 | 92.59 | 7.41 | 88.89 | Down |
| **MCM7** | 24 | 3 | 24 | 3 | 88.89 | 11.11 | 88.89 | Up |
| **MCM3** | 24 | 3 | 24 | 3 | 88.89 | 11.11 | 88.89 | Up |
| **ITGA6** | 24 | 3 | 24 | 3 | 88.89 | 11.11 | 88.89 | Up |
| **SSR2** | 24 | 3 | 23 | 4 | 88.89 | 11.11 | 85.19 | Up |
| **STMN1** | 23 | 4 | 24 | 3 | 85.19 | 14.81 | 88.89 | Up |
| **PRC1** | 24 | 3 | 23 | 4 | 88.89 | 11.11 | 85.19 | Up |
| **POLD1** | 24 | 3 | 23 | 4 | 88.89 | 11.11 | 85.19 | Up |
| **PBK** | 24 | 3 | 24 | 3 | 88.89 | 11.11 | 88.89 | Up |
| **IGSF3** | 22 | 5 | 23 | 4 | 81.48 | 18.52 | 85.19 | Up |
| **DTL** | 24 | 3 | 22 | 5 | 88.89 | 11.11 | 81.48 | Up |
| **ZWINT** | 24 | 3 | 22 | 5 | 88.89 | 11.11 | 81.48 | Up |
| **SPATS2** | 24 | 3 | 24 | 3 | 88.89 | 11.11 | 88.89 | Up |
| **GPSM2** | 23 | 4 | 23 | 4 | 85.19 | 14.81 | 85.19 | Up |
| **COL15A1** | 24 | 3 | 22 | 5 | 88.89 | 11.11 | 81.48 | Up |

Up=Upregulated in cancer or HCC; Down=Downregulated in cancer or HCC; #Up: No of datasets in which gene is over-expressed; #Down: No of datasets in which gene is under-expressed; #Sig: No of datasets in which gene is significantly differentially - expressed; #Non-Sig: No of datasets in which gene is not significantly differentially-expressed; Up (%): Percentage of datasets in which gene is over-expressed; Down (%): Percentage of datasets in which gene is under-expressed; Sig (%): Percentage of datasets in which gene is significantly differentially-expressed

#### 3.2 Gene Enrichment analysis of the Transcriptomic Cores for Hepatocellular Carcinoma

Gene enrichment analysis of these “core genes of HCC” revealed their biological significance as the proteins encoded by the downregulated genes mainly enriched in complement activation and lectin pathways related processes and negatively regulate cellular extravasation. They are also enriched in GO molecular functions like serine-type endopeptidase activity, oxidoreductase activity, RNA methyltransferase activity, etc. (Supplementary Information File 2, Figure S3). Whereas, upregulated core genes enriched in cell cycle GO biological processes like mitotic spindle organization and mitotic sister chromatid segregation, DNA synthesis and DNA replication, postreplication repair and cellular response to DNA damage stimulus, etc. They are also enriched in GO molecular functions such as exodeoxyribonuclease activity, GDP-dissociation inhibitor activity, DNA polymerase activity and insulin-like growth factor binding, etc. (Supplementary Information File 2, Figure S3).

### 3.2 Identification of Robust HCC Biomarkers

With the aim to identify most important genes from the selected set of the signature*, i.e.* “the core genes of Hepatocellular carcinoma”, a simple threshold-based approach (Bhalla et al., 2017; Kaur et al., 2018) and ten-fold cross-validation technique employed using training dataset (GSE25097).

#### 3.2.1 Single Gene Prediction Models using Simple Threshold-based Approach

Here, a simple threshold-based approach employed to distinguish cancer and normal samples. We developed simple threshold-based models for each gene of “the core genes of Hepatocellular carcinoma”. Subsequently, all 26 genes are ranked based on their discriminatory power to distinguish cancer from normal tissue samples as shown in table (Supplementary Information File 1, Table S2). The top 10 genes among the core genes of HCC with the highest performance (i.e. Accuracy > 85%, MCC >0.75 and AUROC >0.85) are selected and enlisted in Table 2.

**Table 2:** Top 10 genes based on the simple threshold-based approach.

| Gene symbol | Thresh | Sens (%) | Spec (%) | Acc (%) | MCC | AUROC | Mean in HCC | Mean in normal | Mean diff |
| --- | --- | --- | --- | --- | --- | --- | --- | --- | --- |
| *FCN2* | 9.78 | 97.76 | 99.59 | 98.63 | 0.97 | 0.98 | 5.76 | 10.89 | -5.13 |
| *CLEC4M* | 7.59 | 97.01 | 98.77 | 97.85 | 0.96 | 0.98 | 4.32 | 9.37 | -5.06 |
| *FCN3* | 10.76 | 95.15 | 99.18 | 97.06 | 0.94 | 0.97 | 7.87 | 12.32 | -4.45 |
| *CLEC1B* | 9.46 | 95.52 | 97.94 | 96.67 | 0.93 | 0.97 | 5.96 | 11.38 | -5.42 |
| *CFP* | 8.14 | 96.64 | 94.24 | 95.5 | 0.91 | 0.96 | 6.15 | 8.63 | -2.48 |
| *CRHBP* | 8.69 | 92.54 | 96.71 | 94.52 | 0.89 | 0.95 | 6.35 | 10.3 | -3.95 |
| *PRC1* | 7.76 | 91.42 | 97.12 | 94.13 | 0.88 | 0.94 | 10.03 | 6.35 | 3.68 |
| *PBK* | 6.03 | 91.04 | 93.42 | 92.17 | 0.84 | 0.93 | 8.65 | 4.41 | 4.24 |
| *DTL* | 6.71 | 85.82 | 94.65 | 90.02 | 0.8 | 0.91 | 8.72 | 5.2 | 3.52 |
| *IGSF3* | 6.93 | 81.34 | 91.77 | 86.3 | 0.73 | 0.88 | 8.1 | 6.08 | 2.01 |

Sens: Sensitivity; Spec: Specificity; Acc: Accuracy; MCC: Mathews Correlation Coefficient; AUROC: Area under Receiver operator curve; Thresh: Threshold; Mean diff: Mean in HCC – Mean in normal

#### 3.2.2 Single Gene Prediction Models using Cross-validation Technique

We have obtained the top 10 genes to classify HCC and normal samples based on simple threshold-based approach. Further, we implemented a ten-fold cross-validation approach to assess the classifying ability of each of these genes using various machine learning classifiers include ExtraTrees, Naive Bayes, KNN, Random forest, Logistic Regression (LR), and SVC-RBF as represented in Table S3 (Supplementary Information File 1). Five out of 10 genes include FCN3, CLEC1B, CLEC4M, PRC1, and PBK are the best performers to distinguish HCC and normal samples with sensitivity, specificity accuracy more than 90%, MCC and AUROC greater than 0.8 and 0.95, respectively as shown in Table S3 (Supplementary Information File 1).

### 3.3 Multiple-Genes based Prediction Models and their Validation on External Validation Datasets

Next, we sought to identify a minimum predictive gene signature that can classify HCC and non-tumorous samples with high precision. Thus, prediction models developed for each gene from the top five genes using different machine learning algorithms like ETREES, Naive Bayes, KNN, Random Forest, LR, and SVC-RBF. Prediction performance of these models was evaluated by using ten-fold cross-validation (CV) on the training dataset. Although models based on each of five genes can distinguish HCC and normal samples of training datasets with high precision, *i.e.* more than 90% sensitivity and specificity. To reduce bias through overfitting and to assure their independence on the testing set, their performance is further evaluated on three external validation datasets. There was a substantial decrease in specificity on these validation datasets (Supplementary Information File 1, Table S4). Therefore, we further developed several models on combining these five genes in various combinations using different machine learning algorithms.

#### 3.3.1.1 The Diagnostic Performance of Five-genes HCC Biomarker-based Models in Identifying HCC

At first all the five genes are combined and subsequently, prediction models developed using different algorithms, *i.e.* ETREES, Naive Bayes, KNN, Random Forest, LR, and SVC-RBF. These models discriminate HCC and normal samples with sensitivity, specificity in a range of 93-97%, an accuracy 95-98%, along with AUROC 0.98-0.99 of training dataset as well as three different cross-platform datasets (shown in Table 3).

**Table 3:**
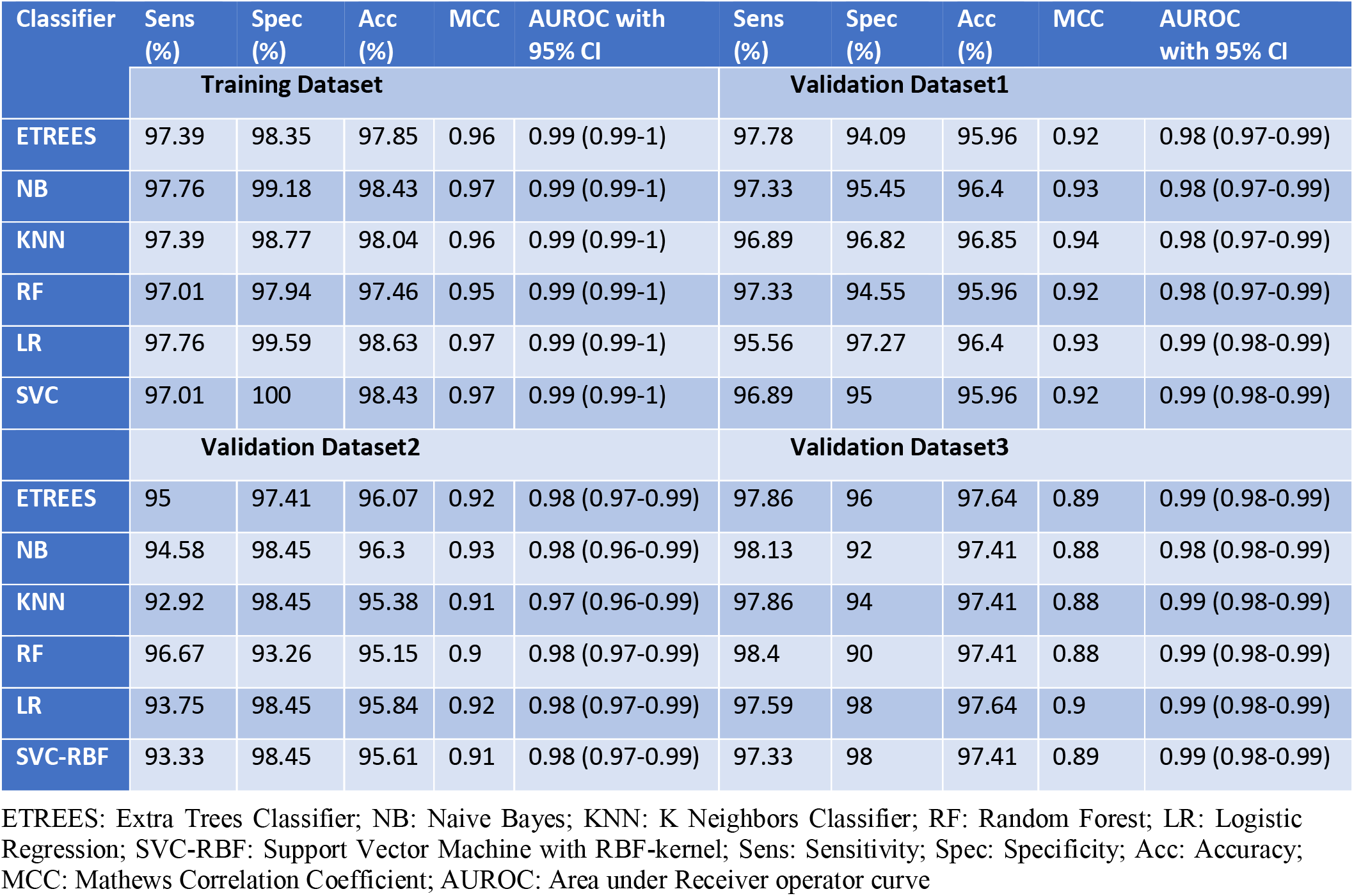
Performance of Five genes (FCN3, CLEC4M, CLEC1B, PRC1, PBK) based models on training and validation datasets implementing various machine learning techniques.

#### 3.3.1.2 Four-genes and Three-genes HCC Biomarker-based Models

To reduce the number of genes from the Five-genes HCC biomarker without compromising the classification performance; we developed several models using the different combination of these five genes, which include Three-genes and Four-genes HCC biomarker sets.

##### 3.3.1.2.1 The Diagnostic Performance of Four-Genes HCC Biomarker based Models in Identifying HCC

Two Four-Genes biomarker panel designed called as Four-genes HCC Biomarker-A contains FCN3, CLEC1B, PRC1, PBK and second Four-genes HCC Biomarker-B contains FCN3, CLEC1B, CLEC4M, PRC1. The prediction models based on Four-Genes HCC Biomarker-A classified HCC and non-tumor samples with good prediction power, i.e. sensitivity and specificity in the range of 93-97% with AUROC in the range of 0.97-0.99 on both training and three independent validation data sets as shown in Table S5 (Supplementary Information File 1).

Prediction models built on Four-Genes HCC Biomarker-B almost performed in similar manner with an accuracy more than 97% and AUROC in the range of 0.97-0.99 on training and two independent validation data sets. While, there is a marginal decrease in the performance on third independent validation dataset, i.e. specificity decreased for Random forest and SVC model in the range 82-92% as shown in Supplementary Table S6.

##### 3.3.1.2.2 The Diagnostic Performance of Three-genes HCC Biomarker based Models for Identification of HCC samples

To further reduces the genes from HCC biomarker panel, two “Three-genes Biomarker” sets designed as Three-genes HCC Biomarker-A contains FCN3, CLEC1B, PRC1 and second Three-genes HCC Biomarker-B includes FCN3, CLEC4M, PRC1. Models based on “Three-genes HCC Biomarker-A” categorize HCC and non-tumor samples very efficiently, i.e. accuracy 95-98% with AUROC in the range of 0.96-0.99 on training as well as independent validation data sets as shown in Table 4. The expression pattern of these three-genes among samples of training dataset and three external validation datasets depicted in Figure 3.

**Table 4:** Performance of Three-genes HCC Biomarker-A (FCN3, CLEC1B, PRC1) based models on training and validation datasets implementing various machine learning techniques.

| Classifier | Sens (%) | Spec (%) | Acc (%) | MCC | AUROC with 95% CI | Sens (%) | Spec (%) | Acc (%) | MCC | AUROC with 95% CI |
| --- | --- | --- | --- | --- | --- | --- | --- | --- | --- | --- |
|  | Training Dataset |  |  |  |  | Validation Dataset1 |  |  |  |  |
| ETREES | 96.64 | 97.94 | 97.26 | 0.95 | 0.99 (0.98-0.99) | 94.67 | 95.91 | 95.28 | 0.91 | 0.97 (0.96-0.99) |
| NB | 97.39 | 99.18 | 98.24 | 0.96 | 0.99 (0.99-1.0) | 96 | 95.91 | 95.96 | 0.92 | 0.98 (0.97-0.99) |
| KNN | 97.76 | 98.77 | 98.24 | 0.96 | 0.99 (0.99-1.0) | 93.78 | 97.73 | 95.73 | 0.92 | 0.97 (0.96-0.99) |
| RF | 97.01 | 97.53 | 97.26 | 0.95 | 0.99 (0.99-1.0) | 94.67 | 96.36 | 95.51 | 0.91 | 0.97 (0.96-0.99) |
| LR | 93.28 | 100 | 96.48 | 0.93 | 0.99 (0.99-1.0) | 92.89 | 97.73 | 95.28 | 0.91 | 0.98 (0.97-0.99) |
| SVC-RBF | 94.03 | 100 | 96.87 | 0.94 | 0.99 (0.98-0.99) | 96 | 96.82 | 96.4 | 0.93 | 0.98 (0.97-0.99) |
|  | Validation Dataset2 |  |  |  |  | Validation Dataset3 |  |  |  |  |
| ETREES | 93.75 | 96.37 | 94.92 | 0.9 | 0.98 (0.97-0.99) | 95.72 | 98 | 95.99 | 0.84 | 0.99 (0.98-0.99) |
| NB | 94.58 | 98.45 | 96.3 | 0.93 | 0.98 (0.97-0.99) | 98.13 | 82 | 96.23 | 0.82 | 0.96 (0.95-0.98) |
| KNN | 95.83 | 97.93 | 96.77 | 0.94 | 0.98 (0.97-0.99) | 97.59 | 96 | 97.41 | 0.88 | 0.99 (0.98-0.99) |
| RF | 95.42 | 94.3 | 94.92 | 0.9 | 0.98 (0.97-0.99) | 95.45 | 96 | 95.52 | 0.82 | 0.98 (0.97-0.99) |
| LR | 95.42 | 98.45 | 96.77 | 0.94 | 0.99 (0.98-0.99) | 97.33 | 98 | 97.41 | 0.89 | 0.99 (0.98-0.99) |
| SVC-RBF | 93.33 | 97.93 | 95.38 | 0.91 | 0.98 (0.97-0.99) | 96.79 | 98 | 96.93 | 0.87 | 0.99 (0.98-0.99) |

ETREES: Extra Trees Classifier; NB:Naive Bayes; KNN: K Neighbors Classifier; RF: Random Forest; LR: Logistic Regression; SVC-RBF: Support Vector Machine with RBF kernel; Sens: Sensitivity; Spec: Specificity; Acc: Accuracy; MCC: Mathews Correlation Coefficient; AUROC: Area under Receiver operator curve.

**Figure 3:**
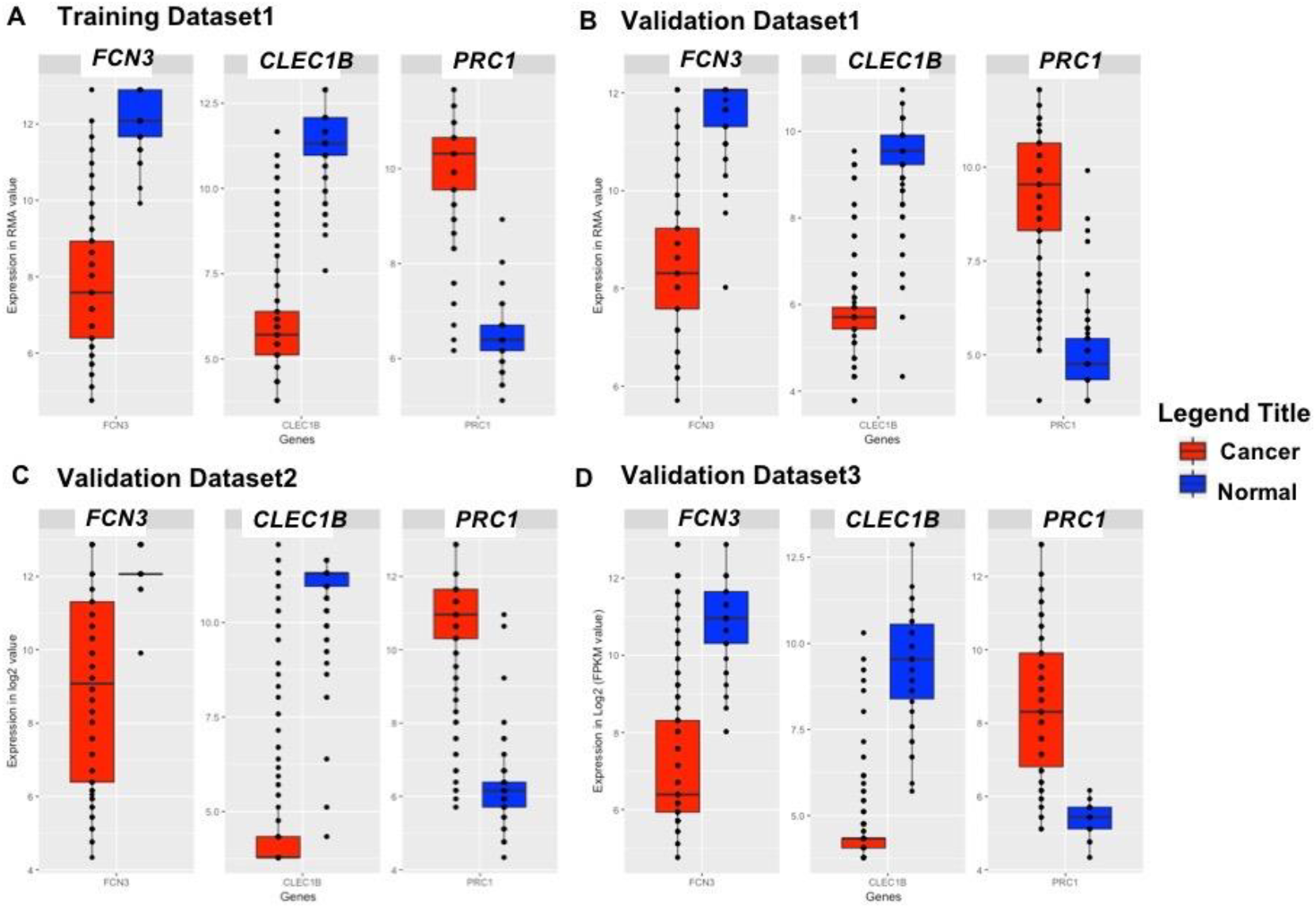
Boxplot representing the Expression pattern of Three-genes panel-based HCC Biomarker.

Most of the models based on “Three-genes HCC Biomarker-B” performed in similar range in classifying samples with a marginal decrease in performance, *i.e.* sensitivity and specificity in range of 91-97% with an accuracy of 92-98% on training and validation data sets as shown in Table S7 (Supplementary Information File 1).

### 3.4 Prediction Model based on the Gene Expression of already identified protein-based biomarker for diagnosis of HCC

To understand the importance of our “Three-genes HCC biomarker”, we also developed prediction models using gene expression of already identified proteins biomarkers, i.e. AFP+GPC3 and AFP+GPC3+KRT19 (CK19) [49]. As we do not have their protein expression for these patients’ samples, hence we employed only their gene expression values. Models based on AFP+GPC3+KRT19 classified HCC and normal samples of training dataset with an accuracy 67-75%. While, this model attained accuracy of 69-77%, 55-87% and 50-74% on external validation dataset1, dataset2 and dataset3, respectively as shown in Table S8 (Supplementary Information File 1). Further, the prediction models based on AFP+GPC3 have improved performance on training dataset with an accuracy of 70-77%, but lower performance on all three validation datasets as given in Table S9 (Supplementary Information File 1).

### 3.5 Prognostic Potential of the “Three-genes HCC biomarker” in the Survival of HCC Patients

To examine the prognostic potential of “Three-genes HCC Biomarker-A”, univariate survival analysis was performed on TCGA and GSE14520 cohorts. The samples were partitioned into low-risk and high-risk groups. Interestingly, all three genes of “Three-genes HCC Biomarker-A” are significantly associated with the survival of HCC. For instance, higher expression (greater than mean) of CLEC1B and FCN3 significantly associated with good outcome of the patients, i.e. OS, DSS, DFS and PFS; while the overexpression of PRC1 is significantly associated with poor survival include DSS, DFS or RFS and PFS of HCC patients for TCGA dataset as shown in Figure 4. In GSE14520 dataset, higher expression of PRC1 is significantly associated with the poor outcome of patients i.e. OS and DFS or RFS, while the higher expression of FCN3 significantly associated with the better outcome of HCC patients as depicted in Figure 5. Complete results of survival analysis with HR (Hazard Ratio), with 95% CI and p-value are represented in Table S10 (Supplementary Information File 1).

**Figure 4:**
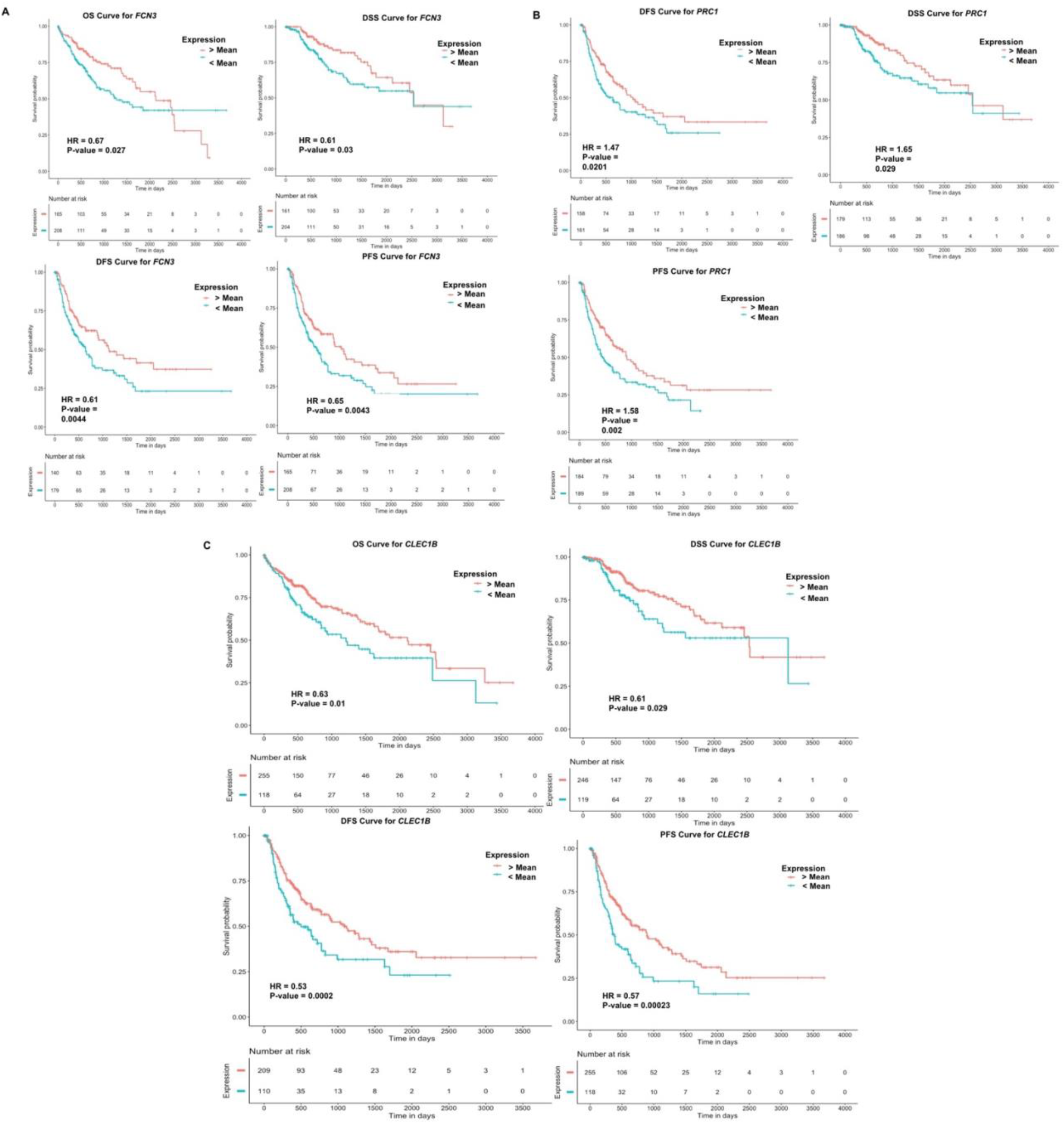
Kaplan Meier survival curves for the risk estimation of HCC patient in TCGA cohort based on the RNA expression of A) *FCN3*, B) *PRC1*, and C) *CLEC1B*

**Figure 5:**
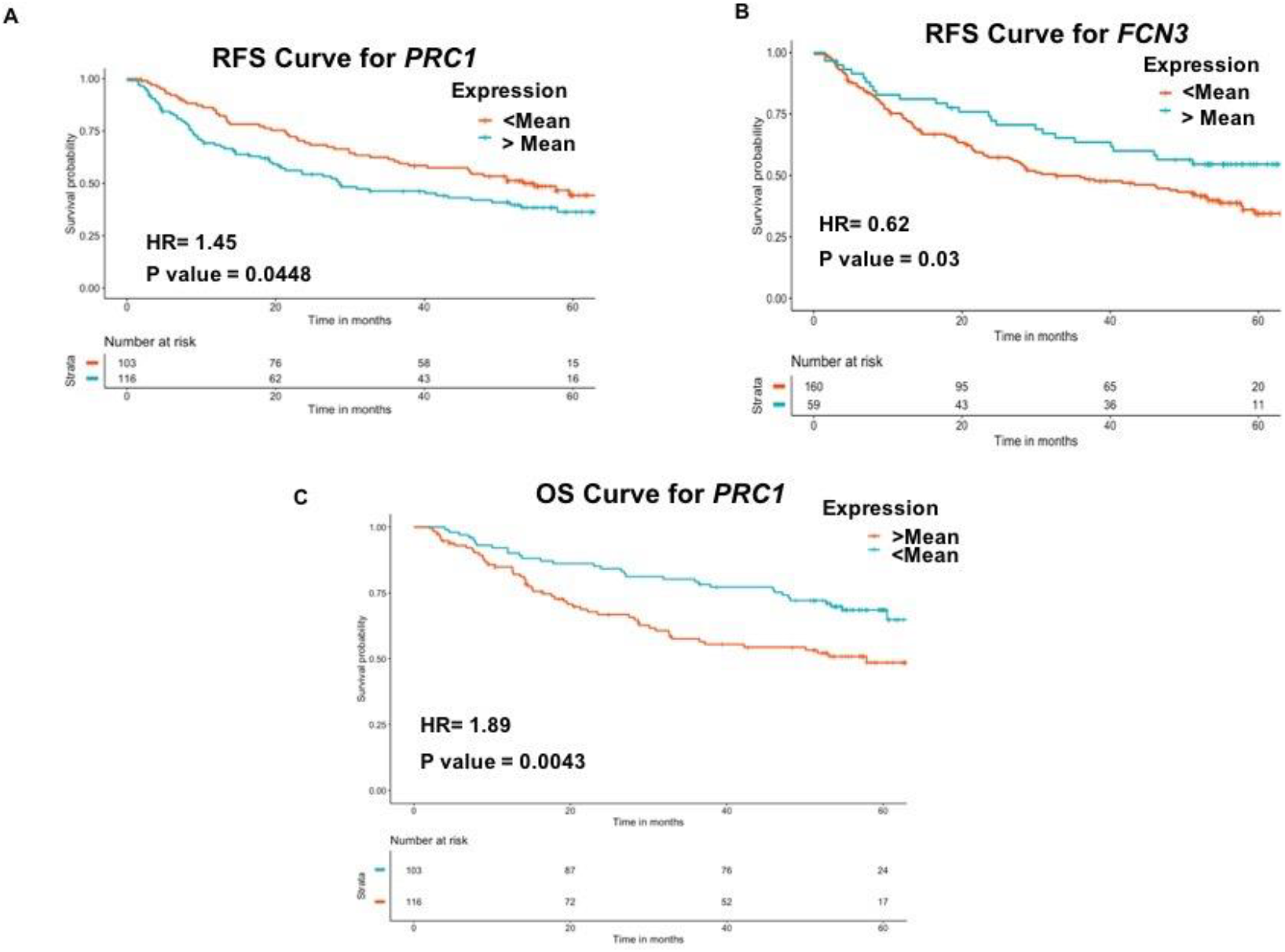
Kaplan Meier survival curves for the risk estimation of HCC patient in GSE14520 cohort based on the RNA expression of A) RFS for *PRC1*, B) RFS for *FCN3*, and C) OS for *PRC1*

### 3.6 Web server

To facilitates the scientific community working in the area of the liver cancer research, we developed HCCPred (Prediction Server for Hepatocellular Carcinoma). In HCCPred, we execute mainly two modules; Prediction Module and Analysis Module based on robust Five-genes, Four-genes and Three-genes HCC biomarkers and 26 Core genes of HCC identified in the present study for the prediction and analysis of samples from the RNA-expression data. The Prediction Module permits the users to predict the disease status, i.e. cancerous or normal using RNA expression values of a subset of genes using in-silico prediction models based on robust Five-genes, Four-genes and Three-genes HCC biomarkers identified in the present study. Here, the user required to submit RMA (for Affymetrix), A-value (for Agilent), Log2 value (for Illumina) or FPKM (High throughput RNA-seq data) for subset of genes or biomarkers. The output result displays a list for patient samples and corresponding predicted status of samples. Moreover, the user can select among the models, i.e. ETREES-based or SVC-RBF based model. Further, the Analysis Module permits the user to analyse the expression pattern of any of top10 ranked genes to check whether it is upregulated or downregulated in comparison to HCC samples based on the samples of the current study. This webserver is freely accessible at webs http://webs.iiitd.edu.in/raghava/hccpred/.

## 4. Discussion

HCC is a type of tumor that is associated with poor prognosis and high mortality rate among all most common cancer types (Siegel et al., 2019). High recurrence rate and low rate of early detection, results in poor prognosis. Accurate diagnosis of HCC may provide the opportunity for appropriate treatment, including traditional available treatment like liver transplantation resection, etc. Although the AFP and DCP proteins are well-established markers for the diagnosis of HCC, but their sensitivity and specificity are not optimum (Sauzay et al., 2016). Therefore, the development of novel robust diagnostic, prognostic biomarker for HCC is needed as it can assist the existing clinical management of tumor.

In this study, we provide a novel large-scale analysis-based approach to identify a robust gene expression-based candidate diagnostic biomarker for HCC derived from multiple transcriptomic profiles/datasets across a variety of platforms obtained from GEO, TCGA. This metadata integration approach across multiple transcriptome datasets of HCC to elucidate core HCC DEG subset followed by class prediction analysis implementing various machine learning algorithms and validation on external independent datasets led us to the identification of multiple-genes based robust biomarkers for HCC. We have identified 26 genes named as “Core genes for HCC” that are uniformly differentially expressed among 80% of datasets. Considering only these genes for downstream machine learning analysis, in an urge to identify a manageable subset with the minimum number of genes from this list that have a high discriminatory power; we further identified “Three-genes based HCC biomarker” which have predictive accuracy of 95-98% and AUROC 0.96-0.99 on training and all three independent validation datasets. Besides, we also developed the prediction models based on the gene expression of already well established protein biomarkers of HCC in the literature, *i.e. AFP+GPC3* and *AFP+GPC3+KRT19* (Lou et al., 2017). The prediction models based on *AFP+GPC3+KRT19* discriminate samples of training dataset with an accuracy of 67-75% and 69-77% of validation dataset1, 55-87% of validation dataset2 and 50-74% of validation dataset3; while the models based on *AFP+GPC3* have quite lower performance on validation datasets. Based on these observations, we hypothesized that “Three-genes HCC Biomarker” might be proved as quite effective diagnostic biomarker for HCC; further protein products of these genes can be also explored from blood or urine, etc. to identify novel protein based non-invasive biomarker as they have very good discriminatory power to distinguish HCC and non-tumor samples at gene expression level from the tissue samples. Moreover, the product of *FCN3* gene is released in the serum and bile (Akaiwa et al., 1999; FCN3 ficolin 3 [Homo sapiens (human)] - Gene - NCBI; Pan et al., 2015; Tizzot et al., 2018), thus this may serve as non-invasive biomarkers for diagnosis of HCC.

Interestingly, the robust “Three-genes HCC Biomarker” contains *FCN3, PRC1, CLEC1B* which have very high diagnostic potential, also possess prognostic potential, *i.e.* they are significantly associated with survival of HCC patients. For instance, higher expression of *CLEC1B* and *FCN3* significantly associated with good outcome of HCC patients in TCGA cohort; while higher expression of *PRC1* is significantly associated with poor outcome of HCC patients in both TCGA and GSE14520 cohorts. Besides, role of *CLEC1B* and *PRC1* was previously also revealed in diagnosis and prognosis of HCC (Chan et al., 2018; Chen et al., 2016; Hu et al., 2018; Kaur et al., 2018).

Taken together, we have established robust three-gene HCC diagnostic biomarker with reasonable performance, possess both diagnostic and prognostic potential; and a meta-data integration pipeline for identification of robust biomarker using machine learning technique, which can work across the different platforms. Further, this pipeline can be used for the analysis of any other cancer type. Although more and more research is under the development of novel biomarkers, further work will be required to implement the clinical utilization of identified biomarker to meet real-world demand. We are anticipating that the identifying novel cost-efficient biomarker using predictive technology for detection of HCC will be promising.

## 5. Conclusions

This study identified and validated a highly accurate Three-genes HCC biomarker for discriminating HCC and non-tumorous tissue samples, also possess a significant prognostic potential that may facilitate more accurate early diagnosis and risk stratification upon validation in prospective clinical trials. Moreover, the product of FCN3 gene is released in the serum and bile, thus this may serve as non-invasive diagnostic biomarkers. Additionally, the uniform overexpression-pattern of *PRC1* among numerous HCC samples suggest it as a novel potential therapeutic target.

## Author Contributions

H.K. collected the data and created the datasets, H.K. developed classification algorithms, H.K. and A.D. implemented algorithms. H.K. and A.D perform the survival analysis. H.K. and A.D. created the back-end server and front-end user interface. S.B., H.K., and G.P.S.R. analysed the results. H.K., R.K., and A.D. wrote the manuscript. G.P.S.R. conceived and coordinated the project, helped in the interpretation and analysis of data, refined the drafted manuscript and gave complete supervision to the project. All of the authors have read and approved the final manuscript.

## Funding

This research was funded by J. C. Bose National Fellowship (with Grant No. SRP076), Department of Science and Technology (DST), INDIA.

## Acknowledgments

All the authors acknowledge funding agencies J. C. Bose National Fellowship (DST). H.K., and R.K., are thankful to Council of Scientific and Industrial Research (CSIR) and A.D. is thankful to DST INSPIRE for providing fellowships, respectively.

## Conflicts of Interest

The authors declare no financial and non-financial conflict of interest.

## Abbreviations

AUROC: Area under the Receiver Operating Characteristic curve
ETREES: Extra Trees Classifier
SVC-RBF: Support Vector Machine with RBF kernel
TCGA: The Cancer Genome Atlas
KNN: K Neighbors Classifier
HCC: Hepatocellular Carcinoma
MCC: Matthew’s correlation coefficient
LR: Logistic Regression
NB: Naive Bayes
RF: Random Forest

## Supplementary Materials

Supplementary Information File 1: Supplementary Tables.

Supplementary Information File 2: Supplementary Figures.

