## Supplementary Information File 2 for "Identification of Platform-Independent Diagnostic Biomarker Panel for Hepatocellular Carcinoma using Large-scale Transcriptomics Data"

**Supplementary Figures:**

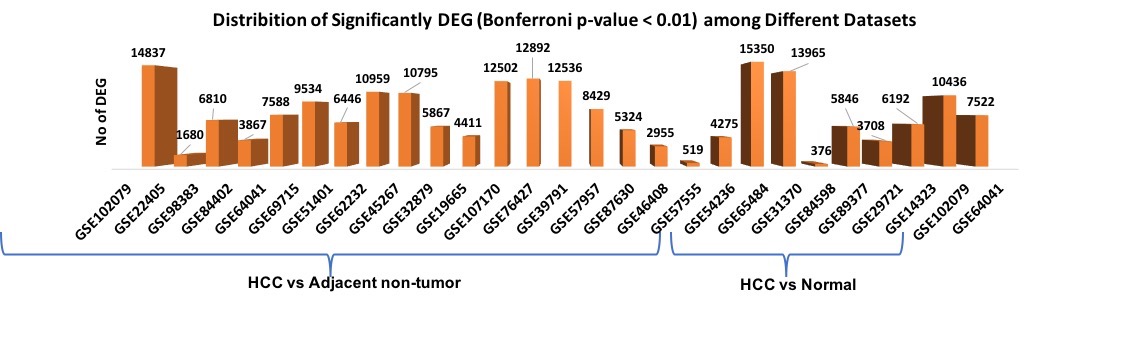

Figure S1**:** Distribution of Significantly DEG (Differentially Expressed Genes) among various datasets with Bonferroni adjusted p-value <0.01.

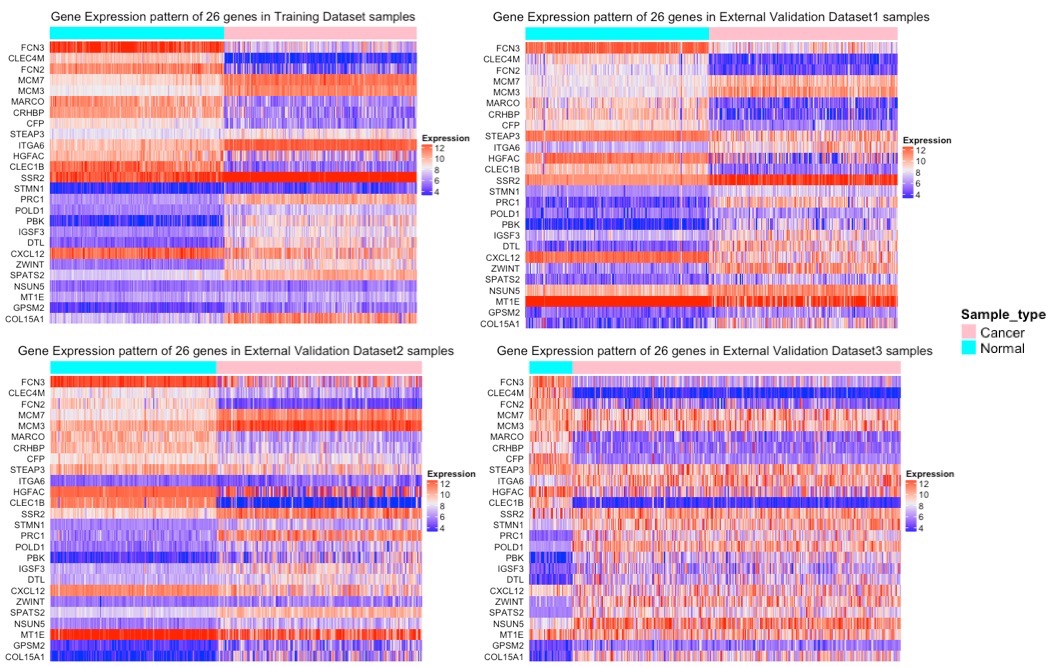

Figure S2: Heatmap representing the expression pattern of “Core genes of HCC” in different datasets.

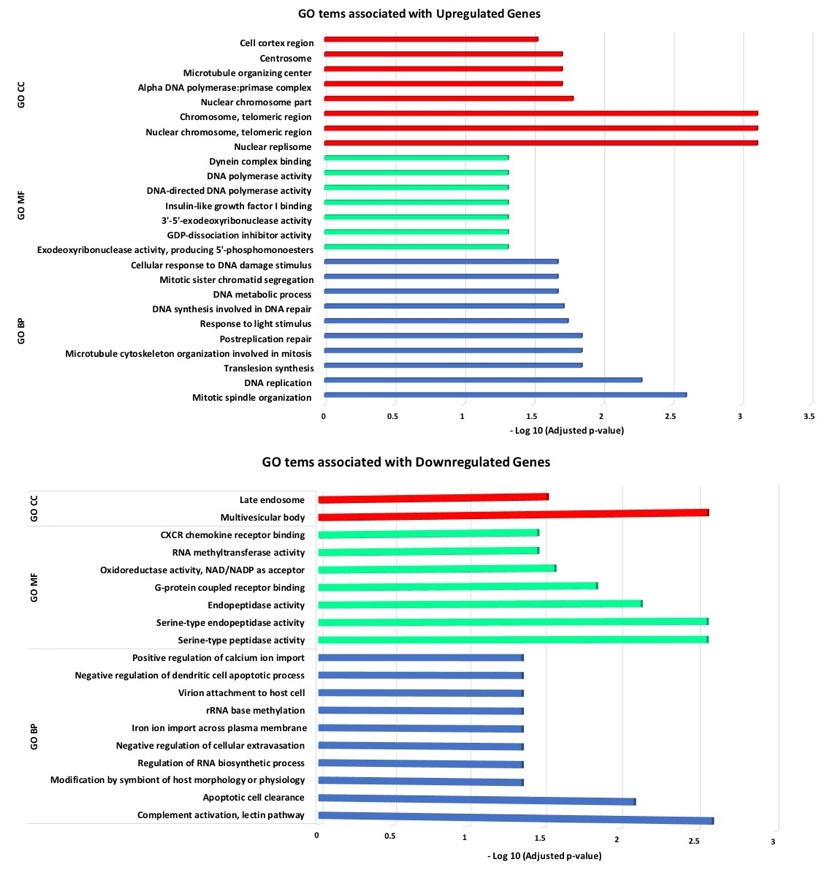

Figure S3: Gene Enrichment analysis of 26 genes or “Core genes of HCC”.
